# Systemic administration of a reported extracellular vesicle inhibitor, dimethyl amiloride, induces preterm birth and fetal growth restriction in pregnant mice

**DOI:** 10.1101/2025.08.13.670125

**Authors:** Madison L. Stone, Scout Bowman-Gibson, Mili S. Bhakta-Yadav, Thomas L. Brown

## Abstract

Successful pregnancy is dependent on extensive coordination between maternal, placental, and fetal units. Alterations in maternal-placental signaling or abnormal placental development can lead to a wide variety of pregnancy complications. This commonly includes preterm birth and fetal growth restriction, which are associated with a high rate of maternal and fetal morbidity and mortality. Maternal-placental signaling is in part mediated by the release of extracellular vesicles. These lipid-bilayer nanoparticles are secreted by various cell types and act as key regulators of both normal and pathogenic cell-cell communication. In a healthy pregnancy, maternal plasma extracellular vesicle concentrations increase as gestation progresses. However, in the pregnancy-associated disorder, preeclampsia, excessive extracellular vesicle secretion occurs and may be involved in the pathogenesis of the condition. Thus, there is a critical need - as a first step - to understand if modifying extracellular vesicle concentration, specifically during pregnancy can attenuate or exacerbate gestational pathogenesis. In this study, we evaluated the effects of administering a reported extracellular vesicle inhibitor, dimethyl amiloride, on maternal extracellular vesicle concentrations in healthy pregnant mice and assessed maternal and fetal outcomes. Maternal administration of dimethyl amiloride resulted in a significant decrease in maternal weight as well as fetal growth and substantially increased the rate of preterm birth. In contrast to previous reports in non-pregnant animals, we found that dimethyl amiloride did not significantly reduce maternal extracellular vesicle concentrations in pregnant mice. Our data demonstrate that systemic administration of dimethyl amiloride drastically impacts the mother and fetus during gestation and caution is suggested against its use during pregnancy.

## INTRODUCTION

Exquisite coordination between maternal, placental, and fetal units is required to achieve a successful pregnancy and is tightly regulated in time and space [1]. Abnormal placental development or alterations in maternal-placental signaling can lead to a wide variety of pregnancy complications including preterm birth and fetal growth restriction which are associated with a high rate of maternal and fetal morbidity and mortality [2,3].

Maternal-placental signaling is in part mediated by the release of extracellular vesicles (EVs) [4–7]. EVs are lipid-bilayer nanoparticles secreted by various cell types throughout the body and act as key regulators of both normal and pathogenic cell-cell communication [8–11]. Upon release into circulation, EVs transport biologically active proteins, miRNA, and DNA cargo to target tissues throughout the body [8–12]. Small EVs (sEVs) are <200 nm and thought to play a central role in cell-cell communication. sEV subpopulations include endosome-derived exosomes (50-150 nm) and small microvesicles (100-200 nm) that directly bleb from the outer cell lipid bilayer [5,10,11,13,14].

Maternal plasma sEV concentrations have been shown to steadily increase as a healthy pregnancy progresses [6,7,15,16]. However, an abnormally elevated level of maternal sEVs has been implicated in the pathogenesis of various pregnancy disorders, particularly preeclampsia [14,17–20]. In non-pregnant animals, systemically reducing elevated sEV levels has been shown to attenuate pathogenic conditions including cancer, chronic kidney disease, and cardiomyocyte hypertrophy [21–24]. However, there is a substantial need to understand whether modifying EV concentration or EV-mediated signaling, specifically during pregnancy, can attenuate or exacerbate gestational pathogeneses as no previous studies have been reported.

5-(N,N-Dimethyl) amiloride hydrochloride (DMA) is a derivative of amiloride. Amiloride is an epithelial sodium channel (ENaC) inhibitor and diuretic used to treat hypertension in non-pregnant individuals [25]. In contrast to amiloride, DMA acts to inhibit Na^+^/H^+^ and Na^+^/Ca^2+^ channels that are present in most cell types, including the placenta [26]. In addition, DMA has been shown to inhibit the transient receptor potential cation channel (TRPA1) which acts as a chemosensor, mechanosensor, and thermosensor in humans and mammals [27,28]. DMA-mediated channel blockade has been reported to inhibit sEV release by reducing intracellular Ca^2+^ concentrations [23,29–32].

In this study, we evaluated the effects of administering a reported EV inhibitor, DMA, on circulating maternal plasma EV concentrations in healthy pregnant mice late in gestation and assessed maternal and fetal outcomes. Many common drugs used to treat non-pregnant individuals are contraindicated in pregnancy due to the potential for fetal growth restriction, preterm birth, fetal demise or fetotoxicity. It is critical to analyze the effects of drugs on maternal and fetal physiology during gestation before exploring their usage in pregnancy-associated disorders.

## MATERIALS AND METHODS

### Vertebrate Animals

All animal procedures were performed in accordance with and approval of the Wright State University Institutional Animal Care and Use Committee (IACUC). Non-pregnant Control (n=5) and non-pregnant DMA-treated mice (n=5) as well as pregnant Control (n=10) and pregnant DMA-treated mice (n=23) were analyzed.

### Isolation of Murine Plasma

Blood was drawn in EDTA-coated tubes (SAI Infusion Technologies, #PMTS-E-1.3), inverted slowly three times and immediately centrifuged for 10 minutes at room temperature at 1,500 x g to obtain platelet-rich plasma. Platelet-rich plasma was subsequently recentrifuged at 4°C for 10 minutes at 2,000 x g to obtain platelet-poor plasma and stored at −80°C until EV isolation and analysis. Plasma with visible hemolysis was excluded from the study.

### Extracellular Vesicle Isolation

EVs from platelet-poor mouse plasma were isolated, purified and analyzed in accordance with guidelines proposed by the International Society for Extracellular Vesicles (ISEV) [13]. Platelet-poor plasma was centrifuged at 14,500 x g for 30 minutes to eliminate cellular debris prior to EV isolation. Exoquick Ultra (Systems Biosciences, #EQULTRA-20A-1) was used to obtain EVs following the manufacturer’s instructions. Briefly, EVs were precipitated for 30 minutes at 4°C and collected by centrifugation at 3,000 x g for 10 minutes. Upon resuspension, EVs were column purified and collected after a final centrifugation at 1,000 x g for 30 seconds. Purified EVs were diluted and analyzed by Nanoparticle Tracking Analysis immediately after isolation.

### Nanoparticle Tracking Analysis (NTA)

EV size and concentration was determined by Nanoparticle Tracking Analysis using a Nanosight NS300 equipped with a 488 nm laser (Malvern Panalytical, software version 3.4). NTA measures the rate of Brownian motion to determine the size and concentration of the nanoparticles [33]. Mouse EV samples were diluted to achieve 20-100 particles per frame.

Samples were analyzed using three 30-second capture videos taken with the following parameters: detection threshold of three and a camera level of 10. Additionally, each sample was recorded with a minimum of 300 tracks per video and analyzed three separate times.

### Western Blotting

Purified EVs were incubated for 15 minutes on ice in 1X RIPA buffer supplemented with protease inhibitor cocktail (Cell Signaling Technology, #9806), sonicated, and frozen at −80°C until use. EV lysates were thawed on ice and 4X reducing sample buffer was added to each sample and heated at 95°C for five minutes. EVs were electrophoresed on 4-15% gradient SDS-polyacrylamide gels. Proteins were subsequently transferred to a methanol-activated Immobilon-P transfer membrane (PVDF, Fisher Scientific) and blocked for one hour at room temperature with blocking buffer [17]. Blots were then incubated overnight with primary antibodies at 4°C for the following proteins: Flotillin 1 (1/1,000 Proteintech, #15571-1-AP), TSG101 (1/1,000 Proteintech, #28283-1-AP), ALIX (1/1,000 Proteintech, #67715-1-Ig), and PLAP (1/1,000 Proteintech, #18507-1-AP). Following secondary antibody incubation at 1/10,000 for 1.5 hours at room temperature, blots were exposed to West Pico SuperSignal enhanced chemiluminescence reagent (Fisher Scientific, #34580), and visualized on an Azure 600 imager (Azure Biosystems).

### *In vivo* administration of DMA

CD-1 mice (8-12 weeks of age, Charles River) were mated overnight. Copulation plug positive females were designated as embryonic day 0.5 (E0.5). At E15.5, pregnant mice were intraperitoneally (i.p.) injected with 7.5 mg/kg DMA (Alomone Labs, #D-165) or 0.9% saline (vehicle Control) for three consecutive days. Maternal weights were recorded daily prior to injection. On E18.5, mice were deeply anesthetized and cardiac puncture was performed to obtain whole blood. Animals were euthanized by CO_2_ followed by cervical dislocation. All procedures were performed in accordance with a Wright State University IACUC approved protocols.

### Statistics

All data presented is reported as mean ± standard deviation (SD). GraphPad Prism 10 software was utilized for statistical analysis and graphing. Statistical significance of the reported data was determined using either two group means by unpaired t-test, or between multiple group means using 2-way Analysis of Variance (ANOVA) with post-hoc Bonferroni correction to evaluate for pairwise comparison. Data with a p-value <0.05 was considered statistically significant.

## RESULTS

### Maternal Analysis

Pregnant mice were treated with DMA on E15.5, E16.5 and E17.5 when EV release is maximal (Fig.1) [34,35]. E15.5 - E17.5 represents a period of rapid fetal growth after placental maturation in mice and coincides with substantial maternal weight gain [1]. To observe the effects of DMA on pregnancy, maternal body weights were recorded daily from E15.5 to E18.5 (Fig.1). Our analysis indicated that daily maternal weight gain was significantly reduced in pregnant DMA-treated females compared to saline-treated pregnant Control mice (Fig.2A). In addition, at E18.5, total maternal weight gain throughout the treatment period was also significantly lower in DMA-treated mice (p=0.0001, Fig.2B).

**Figure 1:**
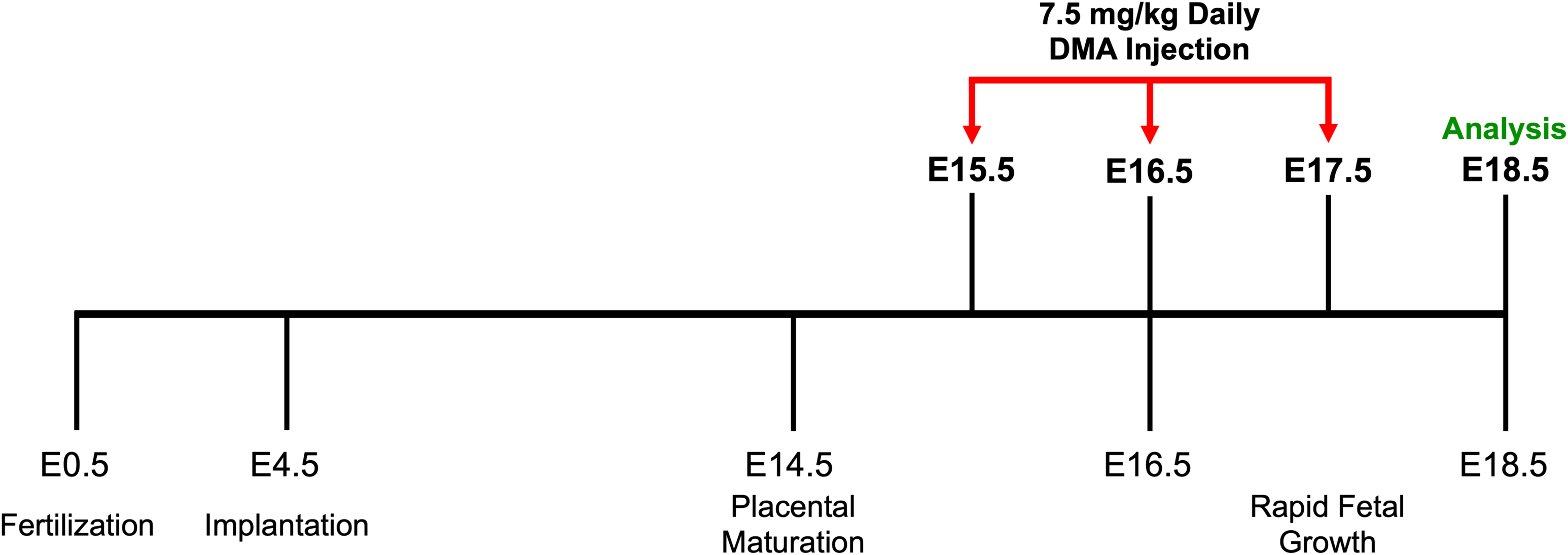
Experimental treatment design. The processes below the timeline indicate important events that occur during normal pregnancy in mice. The events above the timeline indicate the treatment schedule and time of analysis.

**Figure 2:**
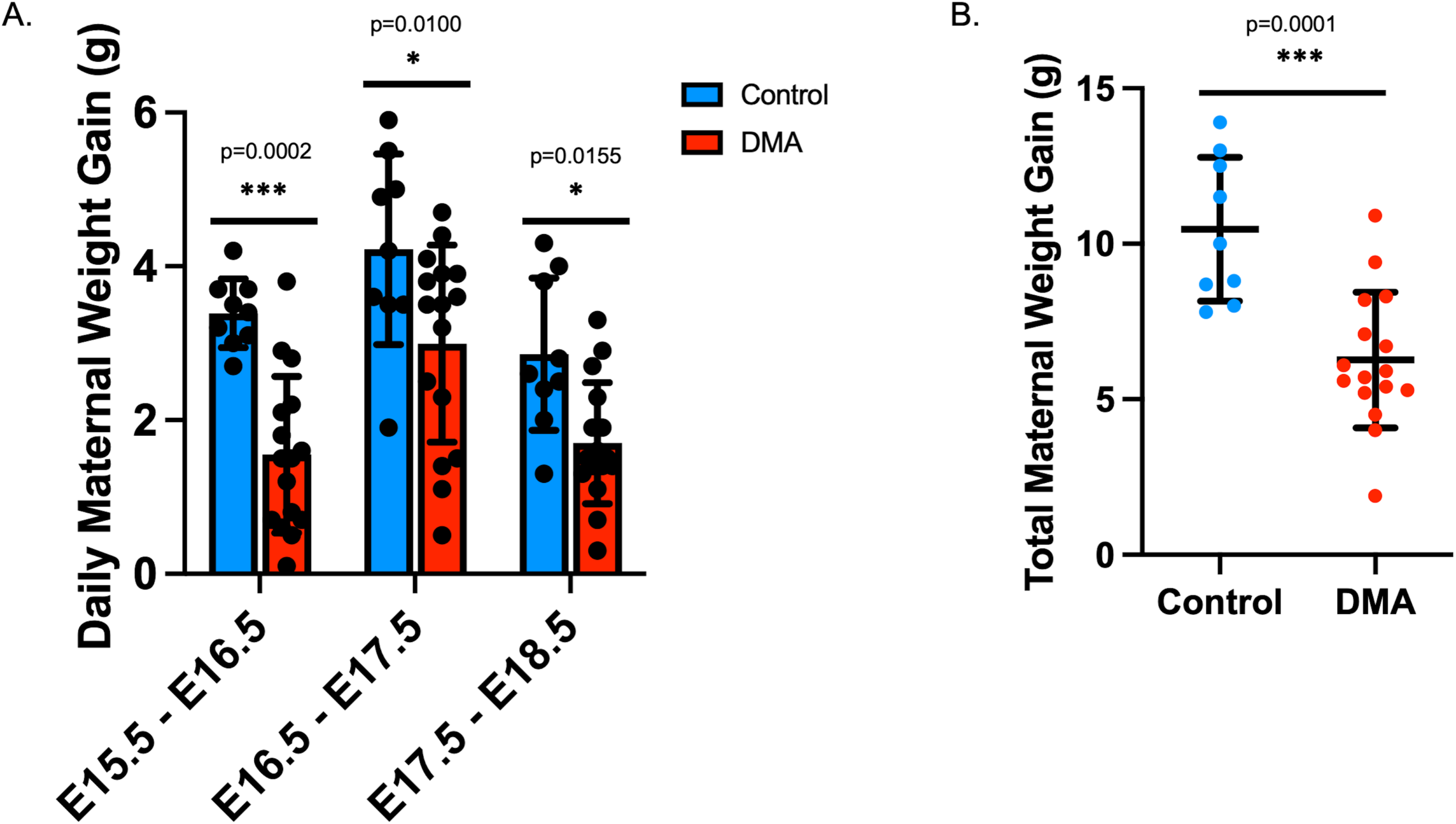
Maternal weight during pregnancy. The amount of maternal weight gained in Control (blue) or DMA-treated (red) pregnant mice. Control (saline) or 7.5 mg/kg DMA were injected on E15.5, E16.5 and E17.5. Maternal weights were recorded 24hrs later - on E16.5, E17.5, and E18.5 - for each pregnancy (2A). The cumulative amount of weight gained throughout the study period in Control (blue) and DMA-treated (red) pregnant mice (2B).

To determine if the effect of DMA on maternal body weight was specific to gestation, we administered either DMA or saline (Control) systemically to non-pregnant female mice for 3 consecutive days (n=5) and monitored their body weight. In contrast to results from pregnant mice, non-pregnant females showed no significant differences in body weight after DMA treatment compared to Controls (Fig.3).

**Figure 3:**
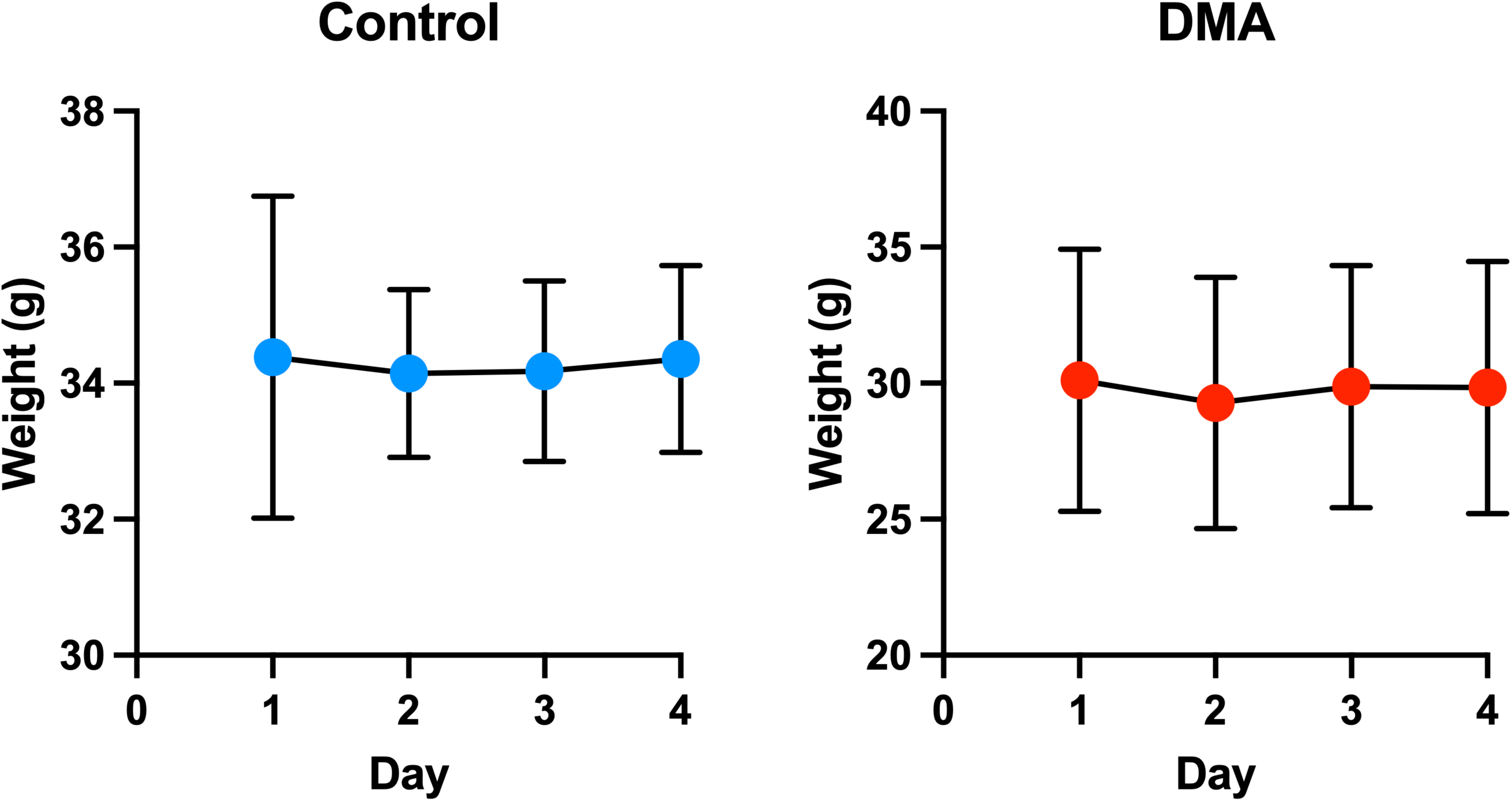
Evaluation of daily weight gain in non-pregnant females. Non-pregnant female mice were treated for three consecutive days with saline (Control, n=5) or 7.5mg/kg DMA (n=5) and daily weights in grams (g) were recorded 24hrs later. All weights were taken prior to injection as well as 24hrs after administration.

### Fetal and Placental Analysis

After three consecutive days of DMA treatment in pregnant females between E15.5 and E17.5, fetuses and placentas were analyzed on E18.5. While normal full-term gestation in CD-1 mice is 19-20 days and spontaneous birth on E19.5 is expected; collection at E18.5 is considered standard for fetal and placental analysis in the field of murine pregnancy. CD-1 mice born before E19.5 are considered developmentally immature and cannot survive outside of the neonatal period [36]. Thus, spontaneous delivery of pups prior to collection on E18.5 was considered premature [36,37]. Our data indicated that preterm birth (PTB) was observed in 30.4% (7/23) of DMA-treated pregnant mice (Fig.4). No PTB was observed in the Control group (Fig.4). Fetuses from DMA-treated pregnancies with PTB were excluded from further investigation.

**Figure 4:**
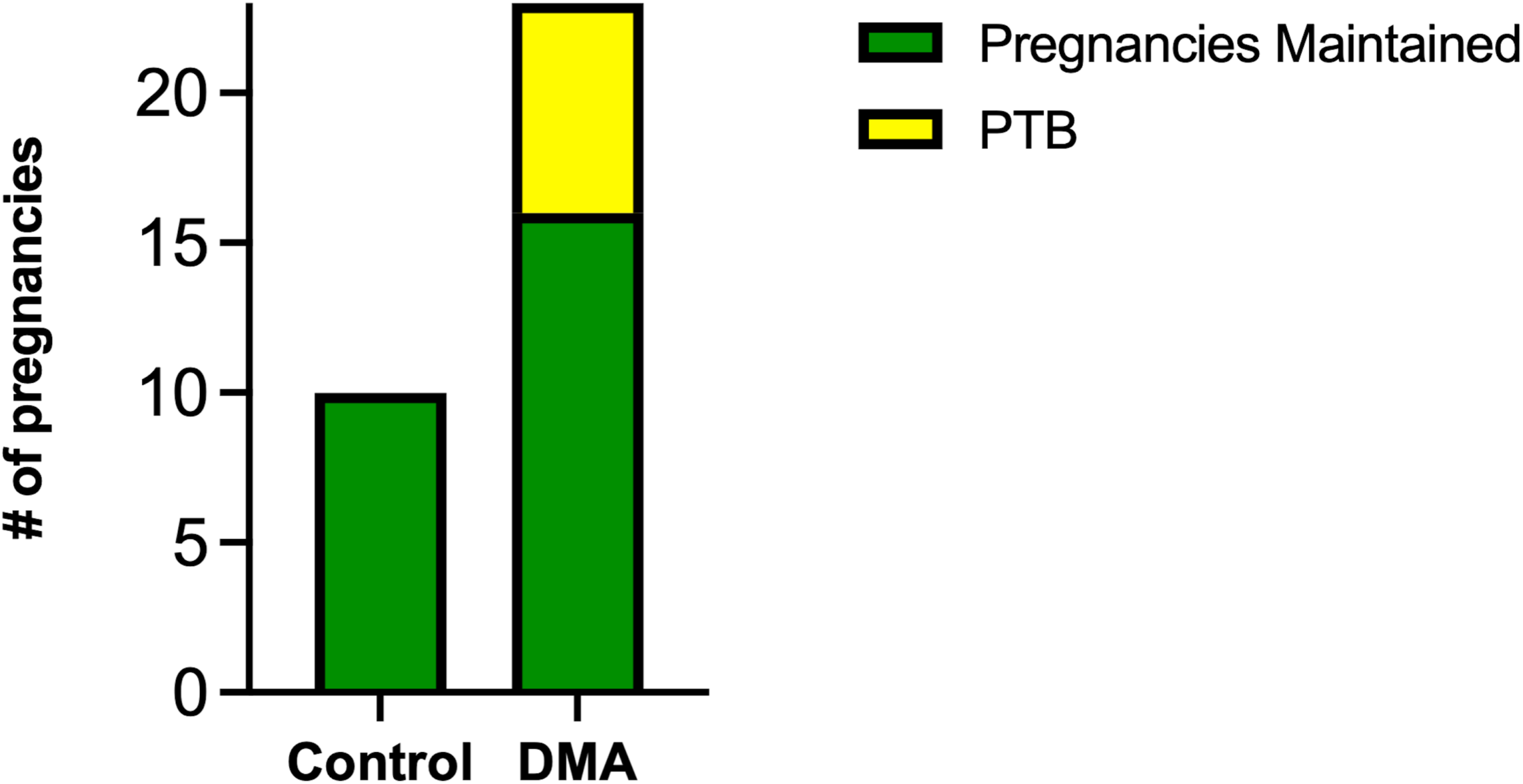
DMA administration induces preterm birth. Full-term gestation in CD-1 mice is 19-20 days and spontaneous birth generally occurs on E19.5. Spontaneous delivery of pups prior to collection on E18.5 is considered preterm. Preterm birth (yellow) was observed in 7 of 23 (30.4%) DMA-treated pregnancies. No preterm birth occurred in the Control group (n=10).

In addition to increased rates of PTB, systemic DMA treatment during pregnancy led to significantly restricted fetal growth (Fig.5A, p=0.0107). Despite this fetal growth restriction, no differences in placental weight or fetal/placental efficiency were observed between pregnant DMA-treated and Control groups (Fig.5B, 5C). In addition, DMA administration during pregnancy did not significantly affect litter size (Fig.5D).

**Figure 5:**
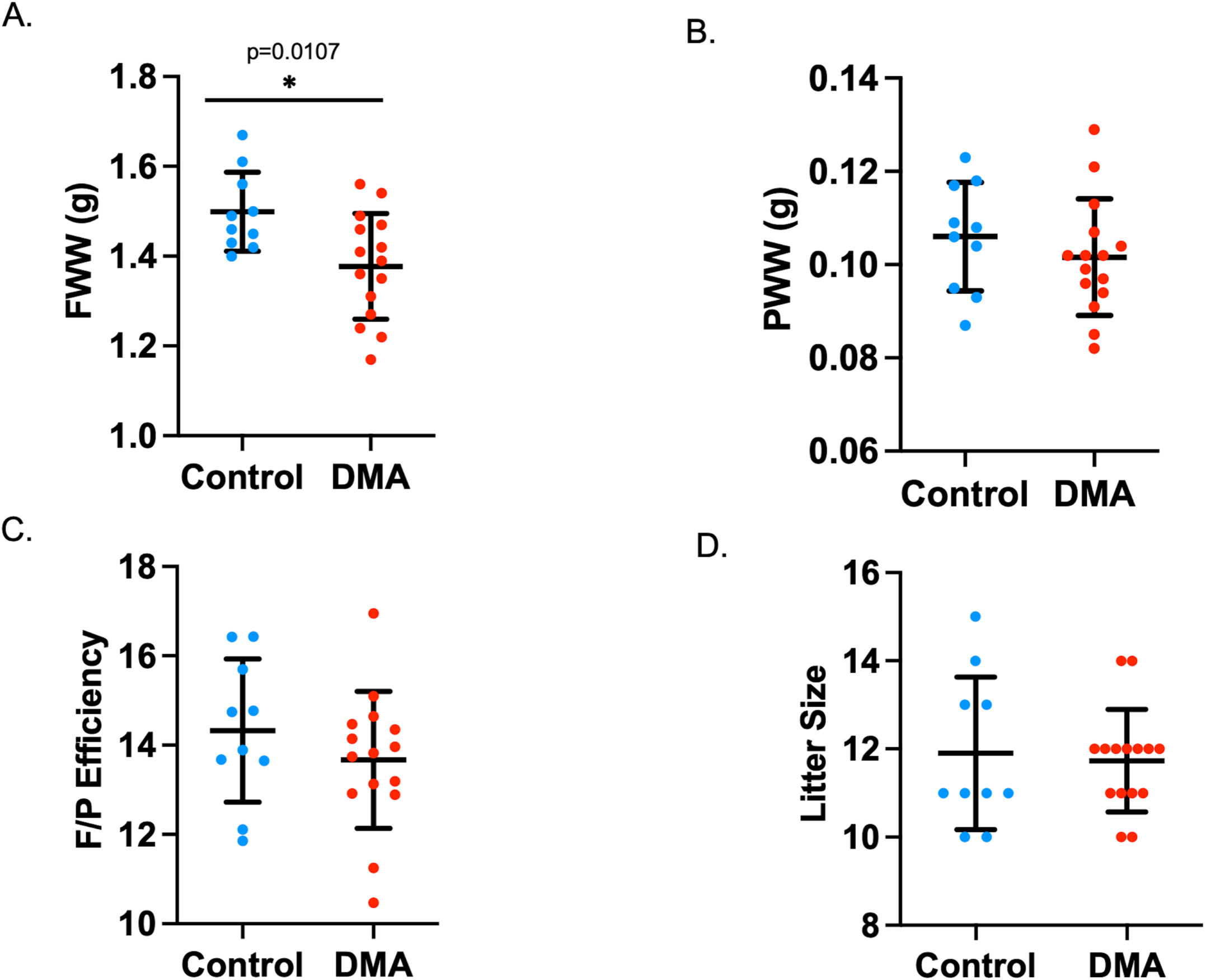
Analysis of fetal and placental parameters. Control (blue) and DMA-treated (red) mice were evaluated for fetal wet weight (FWW, A), placental wet weight (PWW, B), fetal/placental efficiency (C), and litter size (D) for each pregnancy at E18.5. Data points indicate average per pregnancy.

### Extracellular Vesicle Characterization and Quantification

To determine if the fetal growth restriction seen in DMA-treated pregnancies was due to reduced EV secretion, we analyzed EVs in maternal plasma collected from DMA-treated and Control mice at E18.5. Our studies confirmed that analysis of EVs for size and concentration needs to occur immediately after isolation from plasma to prevent freeze thaw effects of particle aggregation and degradation (data not shown) [17,38]. NTA analysis indicated that there was no significant difference in the mean diameter sizes of plasma-derived EVs isolated from pregnant control (111.3 nm +/-10.9 nm) and pregnant DMA-treated (107.7 nm +/-14.1 nm) mice (Fig.6A-C). This size range also confirmed that the purified EVs were representative of small EVs (sEVs) in circulation. Western blot analysis further confirmed the presence of sEVs. Alix and Flotillin 1, markers of exosome biogenesis, were detected in plasma EVs following isolation and purification (Fig.6D). TSG101, a representative protein of exosomal and microvesicular sEV populations, was also present in EVs in both DMA-treated and Control groups (Fig.6D). The detection of placental alkaline phosphatase (PLAP) further indicated that placenta-derived EVs were present within the maternal circulation of pregnant mice, as expected (Fig.6D) [20]. Because no universal marker of EVs has been identified to normalize protein loading, quantification was not determined [13,39].

**Figure 6:**
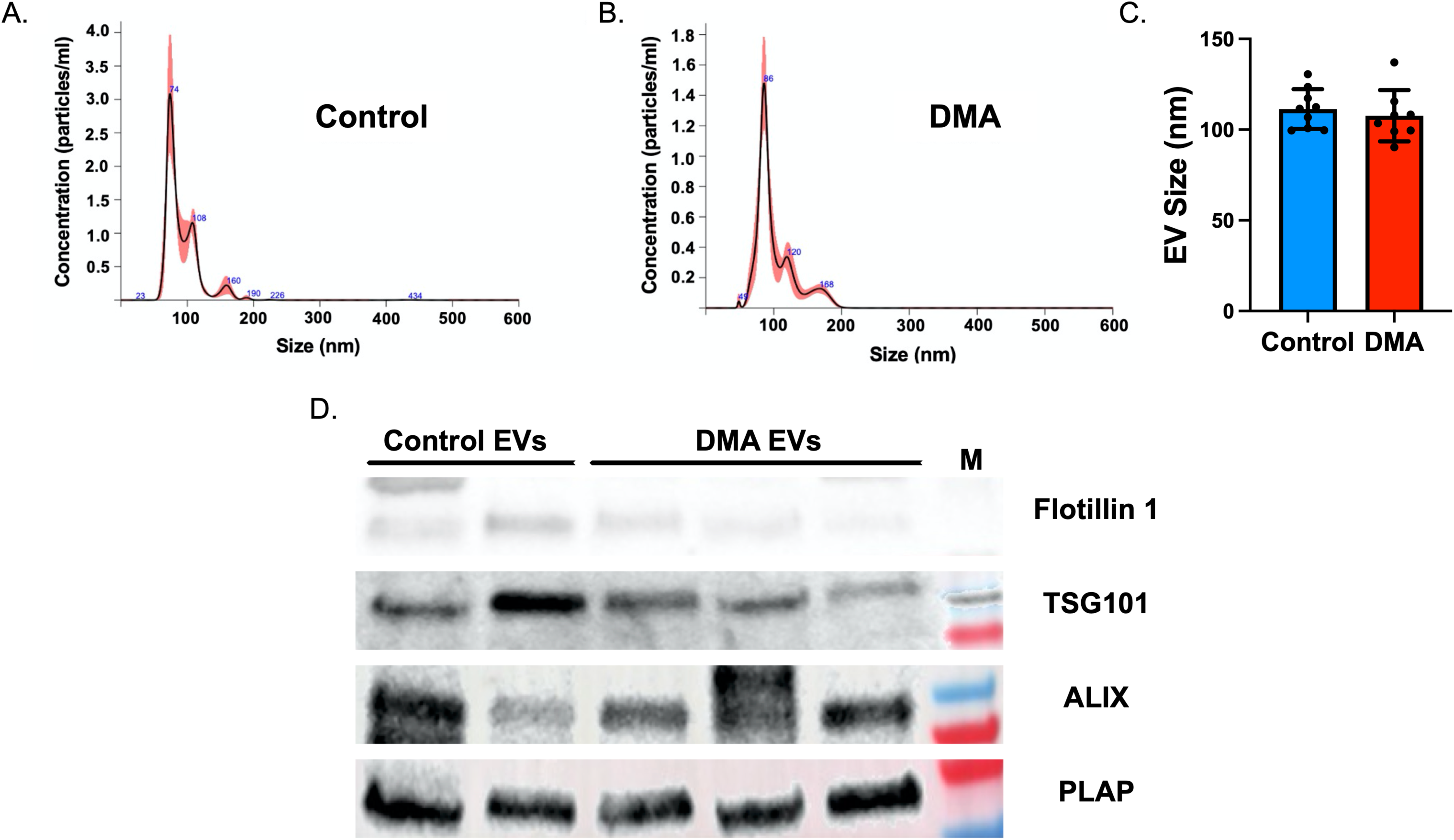
Extracellular vesicle size and analysis. Maternal extracellular vesicle size was determined by Nanoparticle Tracking Analysis immediately after isolation from plasma at E18.5 in Control or DMA-treated pregnant mice. Representative graphs of EV size distributions in Control (A) and DMA-treated mice (B). Analysis of EV size in maternal plasma, per pregnancy, at E18.5 in Control (blue) and DMA-treated (red) mice (C). Western blot analysis of extracellular vesicles isolated from maternal plasma at E18.5 for representative markers of EVs: Flotillin 1 (47kDa), TSG101 (44kDa), ALIX (100kDa) and placental extracellular vesicle marker PLAP (66kDa) in Control and DMA-treated pregnant mice (D). M=Molecular weight marker.

In agreement with previously published human and mouse studies, our data indicate that the concentration of plasma EVs was significantly elevated in pregnant Control mice compared to non-pregnant female Controls (p=0.0450). In addition, EVs were significantly higher in pregnant DMA-treated mice compared to non-pregnant DMA-treated female animals (p=0.0426, Fig.7A) [6,7,15,16,34,35].

**Figure 7:**
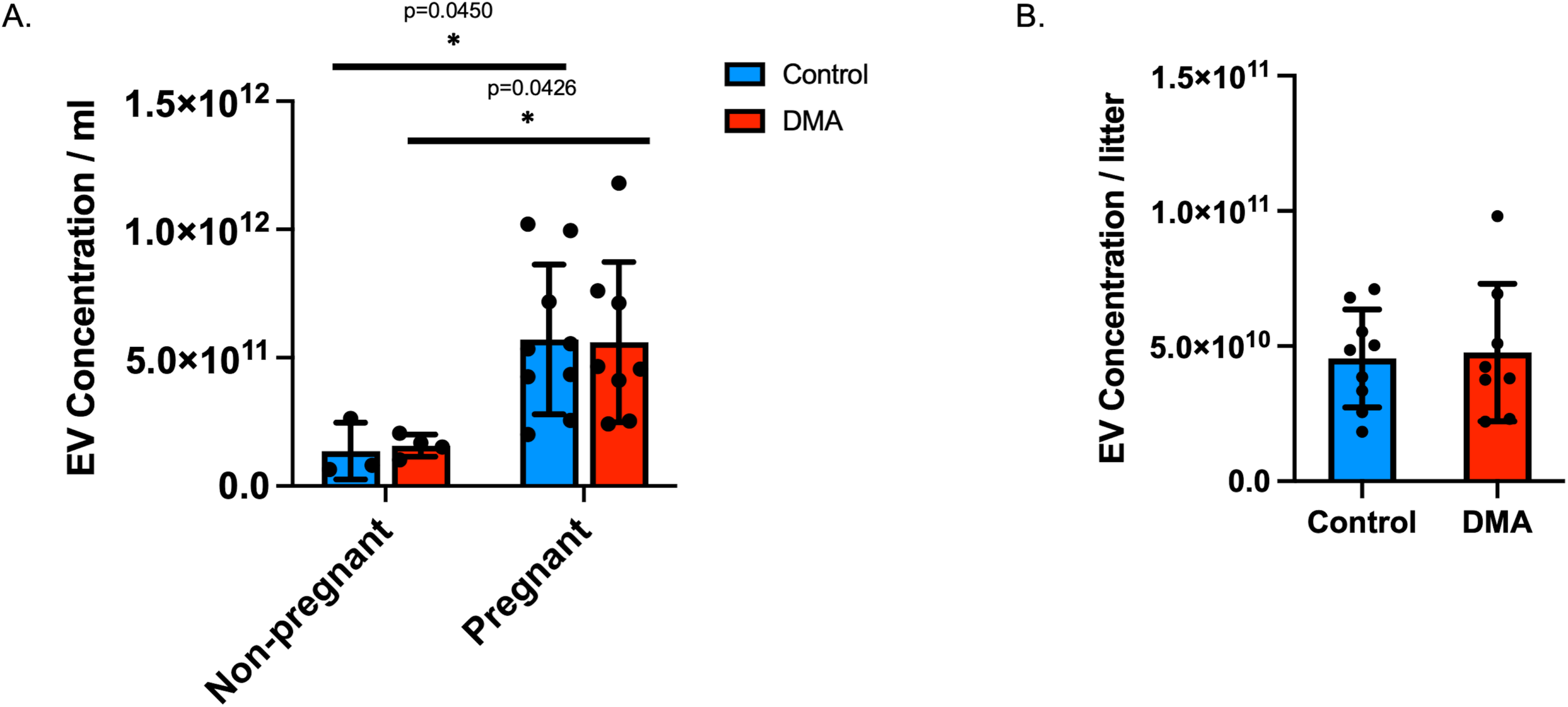
Maternal plasma extracellular vesicle concentrations. Extracellular vesicle concentrations in plasma (particles/ml) were analyzed by Nanoparticle Tracking Analysis in non-pregnant females treated for 3 days with vehicle Control (blue) or DMA (red). In addition, maternal EV concentrations were quantified in pregnant Control (blue) and pregnant DMA-treated (red) females at E18.5 (A). EV concentration was divided by the number of pups to normalize to litter size in Control (blue) and DMA-treated (red) pregnant mice (B).

In contrast to published reports where DMA reduces EV secretion in non-pregnant animals, our data indicate that administration of DMA to pregnant mice from E15.5 to E17.5 did not significantly reduce the concentration of systemic maternal plasma EVs compared to pregnant Control mice (Fig. 7A) [22,23,30,40]. Furthermore, there were no significant differences between DMA-treated and Control groups after maternal EV concentrations were normalized to litter size for each pregnant female (Fig.7B).

## DISCUSSION

The concentration of EVs in maternal circulation is known to increase as a healthy pregnancy progresses [6,7,15,16,34,35]. There is growing evidence to support that abnormally elevated levels of EV secretion may contribute to the pathophysiology of pregnancy-related disorders, particularly preeclampsia [5,14,15,17,18,20]. Patients with preeclampsia, defined as a new-onset hypertension (≥140/90 mmHg) with proteinuria and/or maternal organ dysfunction after 20 weeks of gestation, demonstrate significantly elevated systemic EV concentrations when compared to normotensive pregnant controls [15,17,18,20,41]. It is not yet known if modifying EV secretion during pregnancy could alter the development and progression of preeclampsia. Given that many medications are contraindicated during pregnancy due to their detrimental impact on fetal growth and development, it is important to explore whether EV-targeting treatments are safe to use during gestation.

DMA is a derivative of the diuretic and anti-hypertensive medication, amiloride. DMA is thought to inhibit the release of sEVs by interfering with intracellular calcium accumulation through Na^+^/Ca^2+^ and H^+^/Na^+^ channel blockade [29–31,42,43]. DMA has previously been reported to reduce sEV concentrations and ameliorate fibrotic lesions in injured renal tissues *in vivo* in a male mouse model of chronic kidney disease (CKD), following systemic treatment with DMA (20 mg/kg) for 6 consecutive days [23]. Daily systemic i.p. administration of DMA (∼10 mg/kg) for 7 days has also been reported to decrease sEV release in male mice with pathological cardiac hypertrophy [22]. Daily DMA treatment (∼10 mg/kg) for 7 days, in conjunction with cyclophosphamide, in non-pregnant female tumor-bearing mice has been shown to decrease EV levels and reduce tumor size [30]. While these studies suggest that DMA could reduce sEV secretion for the benefit of specific disease models in non-pregnant animals, evaluation of administering this drug during gestation has never been investigated.

To assess if systemic DMA administration impacts normal pregnancy, we treated healthy pregnant female mice and evaluated the effects on gestational outcomes. Notably, 30.4% of mice treated with DMA delivered preterm, a full 24 hours before normal spontaneous birth is expected to occur. The significantly increased rate of PTB after gestational DMA treatment suggests that DMA alters maternal and/or placental physiology and drastically impacts pregnancy. Because we evaluated mice at E18.5, it is unknown whether more DMA-treated pregnant mice would have spontaneously delivered preterm before E19.5.

In addition to abnormal early spontaneous delivery, maternal weight gain was also significantly reduced from E15.5 to E17.5 in pregnant mice receiving DMA treatment. Conversely, no significant weight changes were observed when DMA was administered to non-pregnant female mice for three consecutive days, indicating that DMA may be eliciting a gestation-specific response that interferes with the normal progression of pregnancy. Numerous complex physiological changes must occur during pregnancy for the maternal/placental unit to adequately support fetal development. These physiologic adaptations could in turn modify the efficacy and activity of DMA, potentially creating a significant threat to maternal and fetal health.

We chose to give daily DMA treatments to pregnant mice from E15.5 through E17.5, as these are the gestational days in which maternal EV concentrations are known to reach maximal levels [34,35]. These days also coincide with a critical gestational period where fetal growth exponentially increases [1]. Importantly, pups evaluated at E18.5 from mice treated with DMA, that did not have PTB, were found to be significantly growth-restricted compared to Controls. While our data indicate a negative effect of DMA on fetal growth, the mechanisms by which this drug resulted in fetal growth restriction are unclear. Nonetheless, the reductions in maternal and fetal weights associated with maternal DMA administration during gestation warrant concern against the use of this drug in pregnant females.

Although DMA-mediated Na^+^/H^+^ and Na^+^/Ca^2+^ channel inactivation has been speculated to reduce sEV secretion in non-pregnant mice, the consequences of blocking the activity of these channels during pregnancy was unclear. To address this, we analyzed maternal plasma sEVs at E18.5 to further evaluate whether the negative pregnancy outcomes seen with DMA treatment were related to reduced maternal plasma sEV concentrations. In contrast to previous reports in non-pregnant mice, our data indicate that no significant reduction in maternal plasma sEVs occurred in pregnant mice after systemic DMA administration. While further investigation would be required to determine specifically how DMA is acting within the maternal and fetal units, the negative outcomes associated with gestational administration of this drug severely limits its usefulness in future studies. DMA is one of several drugs reported to inhibit EV production and/or release; however, most of these compounds have a wide variety of cellular functions and are likely to have significant off-target effects, complicating their use in pregnancy [29,42].

In summary, this study is the first to examine systemically administered DMA to potentially reduce EV concentrations during pregnancy. Our data demonstrate that maternal administration of DMA, late in gestation, resulted in significant decreases in maternal weight as well as fetal growth and a substantially increased the rate of preterm birth. Moreover, DMA did not significantly reduce maternal sEV concentrations when administered to pregnant mice.

While DMA may not affect male or non-pregnant female mice and may reduce EV concentrations when used over a substantial length time in higher doses, our studies demonstrate that DMA drastically impacts pregnancy in mice. Our data provide evidence to caution against systemic administration of DMA as an EV-targeting strategy during gestation.

## ACKNOWLEDGMENTS

This work was supported in part by the Nicholas J Thompson Distinguished Professor Research Grant (TLB), Wright State University Department of Obstetrics and Gynecology (TLB), Wright State University/Premier Health Neuroscience Institute (TLB), and the Wright State University Endowment for Research on Pregnancy Associated Disorders (TLB). We would like to thank the Travers lab at Wright State University for use of the Nanosight instrumentation.

## Declarations

The authors have no competing interests to declare that are relevant to the content of this article. All authors certify that they have no affiliations with or involvement in any organization or entity with any financial interest or non-financial interest in the subject matter or materials discussed in this manuscript.

## Ethics approval

Not applicable.

## Consent to participate

Not applicable.

## Consent for publication

All authors have reviewed and agreed to publication.

## Availability of data and material

All data and material are available upon request.

## Code availability

Not applicable.

